# Genome-wide DNA methylome analysis reveals novel epigenetically dysregulated non-coding RNAs in human breast cancer

**DOI:** 10.1101/002204

**Authors:** Yongsheng Li, Yunpeng Zhang, Shengli Li, Jianping Lu, Juan Chen, Zheng Zhao, Jing Bai, Juan Xu, Xia Li

## Abstract

The development of human breast cancer is driven by changes in the genetic and epigenetic landscape of the cell. Despite growing appreciation of the importance of epigenetics in breast cancers, our knowledge of epigenetic alterations of non-coding RNAs (ncRNAs) in breast cancers remains limited. Here, we explored the epigenetic patterns of ncRNAs in breast cancers via a sequencing-based comparative methylome analysis, mainly focusing on two most popular ncRNA biotypes, long non-coding RNAs (lncRNAs) and miRNAs. Besides global hypomethylation and extensive CpG islands (CGIs) hypermethylation, we observed widely aberrant methylation in the promoters of ncRNAs, which was higher than that of protein-coding genes. Specifically, intergenic ncRNAs were observed to contribute a large slice of the aberrantly methylated ncRNA promoters. Moreover, we summarized five patterns of ncRNA promoter aberrant methylation in the context of genomic CGIs, where aberrant methylation occurred not only on the CGIs, but also flanking regions and CGI sparse promoters. Integration with transcriptional datasets, we found that the ncRNA promoter methylation events were associated with transcriptional changes. Furthermore, a panel of ncRNAs were identified as biomarkers that were able to discriminate between disease phenotypes (AUCs>0.90). Finally, the potential functions for aberrantly methylated ncRNAs were predicted based on similar patterns, adjacency and/or target genes, highlighting that ncRNAs and coding genes coordinately mediated pathways dysregulation in the development and progression of breast cancers. This study presents the aberrant methylation patterns of ncRNAs, which will be a highly valuable resource for investigations at understanding epigenetic regulation of breast cancers.

[Supplemental material is available online at www.genome.org.]

## INTRODUCTION

The development of human breast cancer is driven by both genetic and epigenetic alterations of the cell (Hon et al. 2012; Hervouet et al. 2013). Since the discovery of altered DNA methylation in human cancer, DNA methylation studies in breast cancer have used methodologies of variable scale, focusing on either a few coding genes or regions assumed to be functionally important, such as promoters and CpG islands (CGIs) (Fackler et al. 2011; Hill et al. 2011). However, it is well known that most of the mammalian genome is transcribed producing non-coding RNAs (ncRNAs), the genome-wide methylation patterns of ncRNAs in breast cancers remain largely unknown.

The noncoding RNA transcripts have been classified into a number of subclasses, with the most popular classification being based on their sizes, such as the well-annotated microRNAs (miRNAs) (Osman 2012), long noncoding RNAs (lncRNAs) (Cabili et al. 2011; Derrien et al. 2012). The major class of ncRNAs is lncRNAs, which accounts for about 81.8% of the ncRNAs (Cui et al. 2010). Though the molecular basis of the functions of many lncRNAs are just emerging, many evidences indicate their intricate roles in regulation of a wide variety of biological processes, such as imprinting and gene expression at transcriptional level (Pandey et al. 2008; Ponting et al. 2009; Flintoft 2013). Considering the potential functions of lncRNAs, their transcription must be tightly regulated. Aberrant expression of lncRNAs has appeared in prevalent cancer types, including breast cancer. A notable example is *HOTAIR* that is over-expressed in breast cancers, and loss of *HOTAIR* moderates the invasiveness of breast cancer (Nie et al. 2013). Another example is the lncRNA *MIR31HG*, which is expressed abundantly in the non-invasive breast cancer cell lines of luminal subtype (Augoff et al. 2012). While lncRNAs are a proven component of gene expression modulation (Mercer and Mattick 2013), epigenetic regulation of lncRNA itself remains poorly understood. Emerging studies have described aberrant methylation of specific lncRNAs in breast cancers; studies of the aberrant epigenetic regulation patterns at a global scale of lncRNA genes are scarce.

In addition, it has become apparent that miRNAs have been one of the recently discovered and well characterized classes of ncRNAs (Griffiths-Jones et al. 2006). MiRNAs are important regulators of gene expression that are frequently deregulated in cancers (Wu et al. 2010; Iorio and Croce 2012), with aberrant DNA methylation being an epigenetic mechanism involved in this process (Croce 2009; Suzuki et al. 2012; Baer et al. 2013). Aberrant DNA methylation linked to silencing of individual miRNAs has been demonstrated in many cancer types, including breast cancer (Weber et al. 2007; Shen et al. 2012). Some of these miRNAs function as tumor suppressors (such as *miR-203*, *miR-195* and *miR-497*) and their down regulation due to aberrant hypermethylation is associated with increased malignancy or metastatic potential in breast cancer (Li et al. 2011; Zhang et al. 2011b). Using 5-methylcytosine immunoprecipitation coupled to miRNA tiling microarray hybridization, Vrba et al. have demonstrated that miRNA gene promoters are frequent targets of aberrant DNA methylation in human breast cancer (Vrba et al. 2013), indicating an important role of DNA methylation in miRNA deregulation in cancer. However, only 167 miRNAs were analyzed in their study, accounting for only 10% miRNAs in the genome. To our knowledge, a comprehensive analysis of DNA methylation of miRNAs in breast cancer is still insufficient.

The next-generation sequencing technologies have emerged as powerful tools to allow whole genome profiling of epigenetic modifications, including DNA methylation. MBDCap-seq protocol, for instance, is a technique used to capture methylated DNAs by using a methyl-CpG binding domain (MBD) protein column followed next-generation sequencing (Robinson et al. 2010). The low cost and unbiased display of methylation profiles of both coding and non-coding regions make it suitable for genome-wide methylation profile analysis. Here, we reported use of high throughput sequencing technology to map DNA methylation in a cohort of 87 breast samples (77 cancers and 10 normal controls) (Gu et al. 2013). Comparative analysis of the methylomes presented the unbiased systematic effort to determine the aberrant methylation patterns of ncRNAs, and provided the precise genomic locations that undergo methylation changes, which will be a highly valuable public resource for investigations at understanding epigenetic regulation of the breast cancer genomeand identification of ncRNA therapeutic targets.

## RESULTS

### Global differences of DNA methylation in breast cancer

We performed comprehensive comparison analyses of DNA methylation profiling in normal controls and breast cancers, which were downloaded from the Cancer Methylome System (CMS) (Gu et al. 2013). Pair-wise comparison of average genome-wide methylation intensity across all breast cancers and normal cohorts showed that normal and breast cancer samples correlated reasonably well within classes, but not inter-classes (Figure 1A). This result remained valid even when we used more relaxed resolutions of genomic regions (Figure S1), indicating that a large fraction of the breast cancer genome is differentially methylated as compared to normal cells. Moreover, the correlation coefficient among tumor group shows higher variation than that of the normal group (Figure 1B), suggesting that some heterogeneity might be in the methylation profiles among breast cancers.

**Figure 1.**
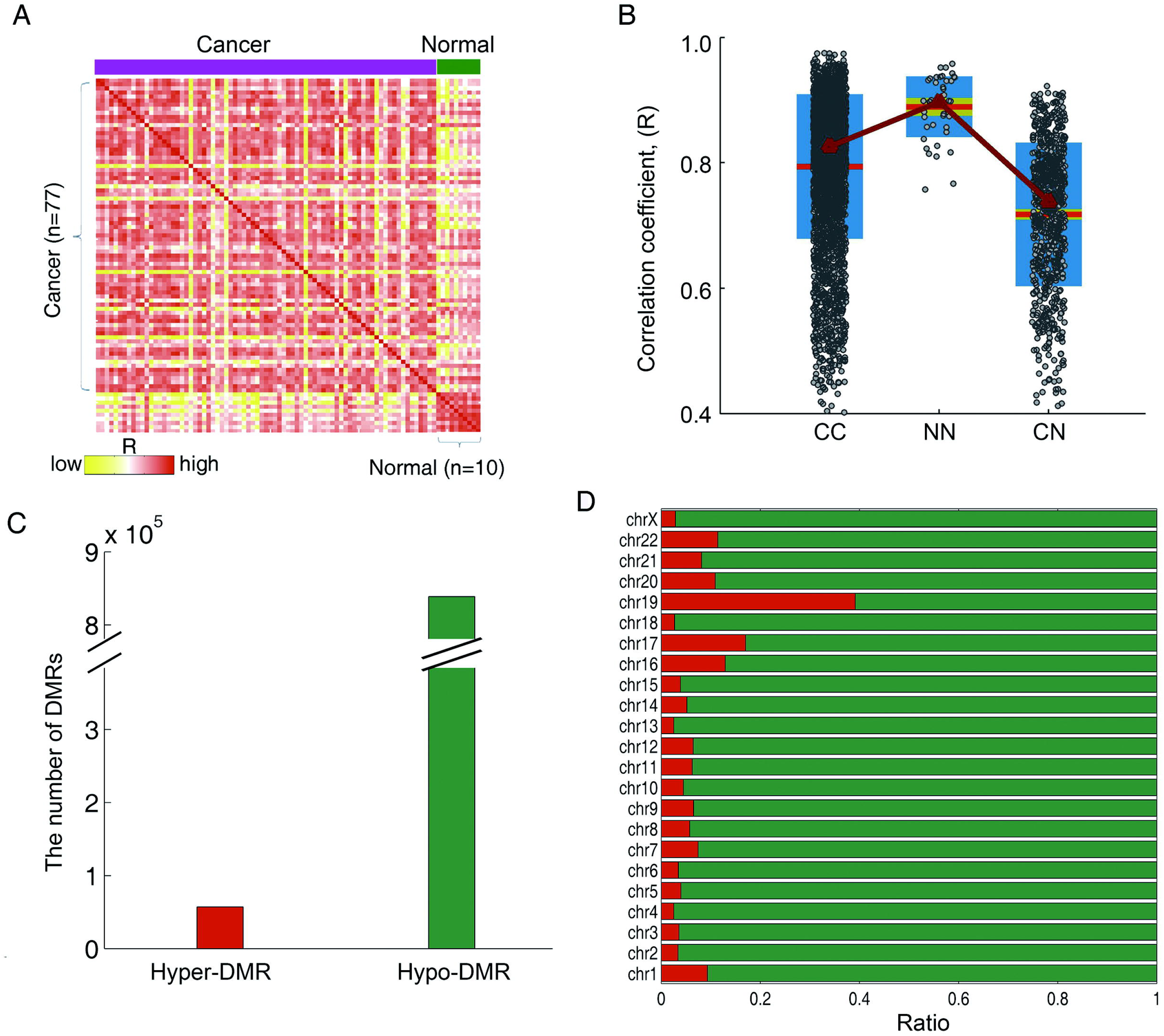
Global DNA methylation profiles highlight clear difference in breast cancers and normal controls. (A) DNA methylation was calculated from MBD-seq data using 1 kb bins across the genome, and the relationship among samples was determined by Pearson correlation. (B) The distribution of Pearson correlation among samples. Red lines indicate the mean correlation coefficient and the light green areas represent of the 95% confidence interval of mean. The dark blue areas indicate the stand error of the correlation coefficient, and the median correlation coefficient is fitted by the dark red line. Each dot represents the correlation between two samples. (C) The number of DMRs identified in the comparison between breast cancer and controls. (D) Stacking barplots showing percentage of hyper and hypomethylated DMRs out of all bins for each chromosome.

Next, we carried out statistical comparisons between the breast cancer methylomes with the controls to identify differentially methylated genomic bins (see details in Methods). As a result, we observed global hypomethylation and local hypermethylation in breast cancer compared to controls. A striking of 4.72% genome was hypomethylated; on the other hand, hypermethylation was relatively limited, with 0.45% genomic bins exhibiting hypermethylation compared to controls. And then consecutive differentially methylation bins with no gap between them were merged into differentially methylation regions (DMRs). As a result, we identified about 838,817 hypo-DMRs in breast cancers, which were about sixteen times of hyper-DMRs (Figure 1C). Investigation of the distribution of hypermethylated and hypomethylated DMRs across the entire genome, we found that the hyper- and/or hypo-methylated DMRs were dispersed throughout the genome on multiple chromosomes and in each chromosome, the proportion of hyper- and hypo-methylated DMRs is similar (Figure 1D). In summary, the comparison analyses of comprehensive methylation profiles indicated that there are extensive methylome alterations in breast cancers.

### Extensive CGIs hyper-methylation in breast cancer

Previous DNA methylation analyses have mainly focused on CpG islands (CGIs), which are highly clustered CpGs often located at gene promoter regions (Hatada et al. 2006; Dai et al. 2013). Mapping the DMRs to CGIs, we observed that most (31.85%) of the CGIs showed hyper-methylation in breast cancers (Figure 2A), associated with ∼25.20% hyper-DMRs, which was significantly higher than random regions (p<0.001, Figure S2A, details in Text S1). Besides CGIs, CpG island shores had been demonstrated to be frequently aberrant methylation in colon cancer (Irizarry et al. 2009). However, investigation of the methylation patterns of CGI shores in breast cancer is hampered by the lack of genome-wide profiling. We found that about 40% of the CGI shores, both 5’-shores and 3’-shores showed significantly aberrant methylation (Figure 2A), wherein 20.11% and 20.76% of 5’-shores and 3’-shores were respectively hypermethylated, covering more than 20% hyper-DMRs, which were significantly higher than random regions (Figure S2B). On the other hand, the hypo-DMRs were significantly under-represented on CGIs and shores (p<1.0e-3). Of the hypermethylated CGIs, 88.69% were also mapped with hyper-DMRs on their shores, either on 5’-shores or 3’-shores (Figure 2B). However, of the hypomethylated CGI shores, few (0.28%) corresponding CGIs were aberrantly methylated. In addition, we found that the 5’-shore and 3’-shore were rarely both hypomethylated (Figure 2C). Analyses of the distribution of aberrant methylation frequency around hyper- and hypo-methylated CGIs, there was a reversal in the frequency trends observed between hypermethylation and hypomethylation (Figures 2D and 2E). The hypermethylation frequency was extremely high in CGIs, while the hypomethylation frequency was substantially high on CGI shores. Collectively, these data indicated that in breast cancer, a large proportion of CGIs were hypermethylated which may drive a specific transcriptional program to promote breast carcinogenesis.

**Figure 2.**
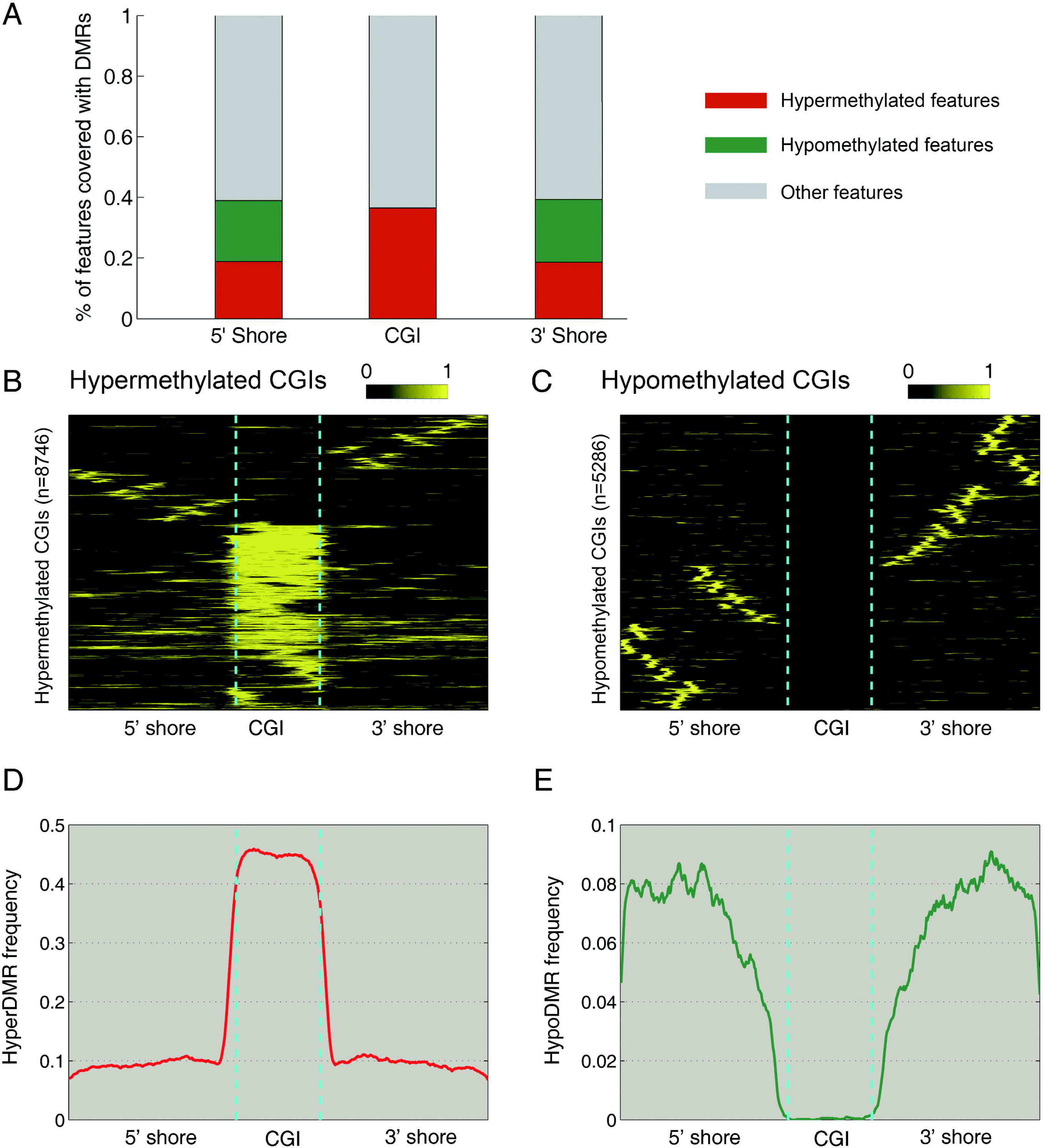
Extensively CGI hypermethylation in breast cancers. (A) Stacking barplots representing the percentage of hyper- and hypomethylated CGIs and shores. (B) and (C) show the aberrant hyper- and hypomethylation around the CGIs. Each row represents a unique CGI and corresponding shores, which was divided into 500 windows. The aberrant methylation frequency is indicated in yellow. (D) and (E) show the average aberrant hyper- and hypo methylation frequency around the CGIs, respectively.

### Aberrant methylation is associated with ncRNAs in breast cancer

Promoter aberrant methylation is usually linked to an altered chromosomal state, and thus to transcriptional gene silencing, which has been thought to contribute to tumorigenesis (Guzman et al. 2012). To determine whether DMRs preferentially occurred in promoters, genome-wide DMRs were mapped to coding and non-coding RNA promoters, exons and other genomic features. As a result, we identified 11.49% promoters of coding genes that were aberrantly hypermethylated in breast cancers (Figures 3A and 3B), associated with 28.53% hyper-DMRs which were significantly higher than random regions (p<0.001, Figure S2). On the other hand, about 19.77% coding gene promoters were hypomethylated in breast cancers, which were associated with about 3.23% hypo-DMRs. Collectively, these results indicated that coding gene promoters were preferentially aberrantly methylated in breast cancers. Among these aberrantly methylated coding gene promoters, several were previously reported to be involved in cancer pathogenesis, such as, *HOXB2*, *FGF4* and *TTK* (Li et al. 2008; Marsit et al. 2010; Maire et al. 2013). Our observations of aberrant methylation of these genes provided a new layer of epigenetic regulation in breast cancers.

**Figure 3.**
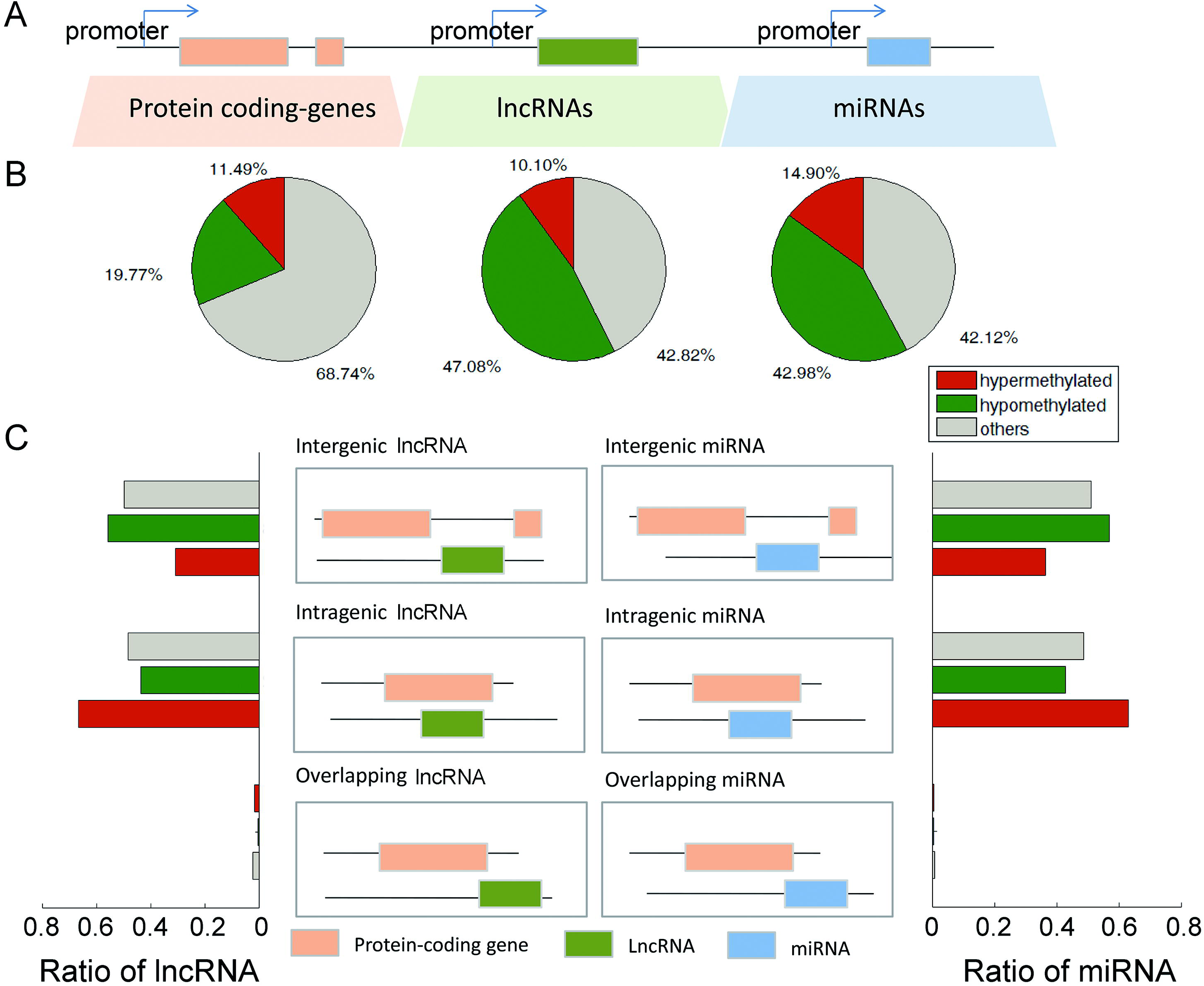
Characterization of genome-wide methylation patterns in ncRNAs. (A) Cartoon represents the genomic features analysed. (B) Pieplots represent the percentage of coding and non-coding promoters overlapping with hyper- and hypo-DMRs in breast cancer. The green pies represent the percentage of features covered with hypo-DMRs and the red ones represent the percentage of hypermethyalted features. (C) LncRNA and miRNAs were classified into different subcategories based on their location with protein coding genes. Barplots show that the majority of intragenic ncRNAs were overrepresented in the hypermethylation and most of intergenic ncRNAs were hypomethylated in breast cancers.

In addition to aberrantly methylated coding genes identified above, non-coding RNAs (miRNAs and lncRNAs), are important regulators of gene expression. Compared with the proportion of aberrantly methylated coding gene promoters, there were more ncRNA promoters that were aberrantly methylated in breast cancers (Figure 3B). Specifically, about 57.18% of lncRNA promoters and 57.88% of the miRNA promoters were aberrantly methylated, respectively (Figure 3B). In addition, 3,644/852 hyper-DMRs and 14,847/2,030 hypo-DMRs were separately mapped to lncRNA and miRNA promoters (Figure S2C). Although a large proportion of coding and non-coding gene promoters were covered with hypomethylated DMRs, no significant overrepresentation were observed comparing with random regions (Figures S2B and S2C). These results suggested that the promoters, especially the ncRNA promoters, undergo preferential aberrant hypermethylation during the breast cancer development.

Previous reports have suggested that lncRNAs can affect the expression of nearby protein-coding genes or act at a distance to control broader biological processes (Ponjavic et al. 2009; Jia et al. 2010). On the other hand, evidence has shown that intronic and intergenic miRNAs were with different numbers of target binding sites (Assel S. Issabekova 2011). These results indicated that different biotypes of ncRNAs were with distinct functions. Thus, these ncRNAs were further reclassified into three biotypes based on their location with respect to protein-coding genes: intergenic, intragenic and overlapping ncRNAs. Next, we investigated the methylation of these ncRNA biotypes. As a result, intergenic lncRNAs were observed to contribute a large slice of the aberrantly methylated lncRNA promoters in breast cancers. These intergenic lncRNAs could not be detected on the microarray previously, and our genome-wide analysis provided new candidates for further functional analyses. Specifically, about 31.79% of the hypermethylated lncRNAs were intergenic lncRNAs and 66.81% of the hypermethylated lncRNAs were intragenic. On the other hand, hypomethylated lncRNAs were overrepresented in the intergenic biotype. MiRNA shared similar patterns with lncRNAs, 62.94% of the significantly hypermethylated miRNAs were intragenic in breast cancers while hypomethylated lncRNAs were overrepresented in the intergenic subtypes (Figure 3C). These results indicated that the aberrantly methylated lncRNAs may share the common patterns with their host genes. Indeed, about 79.86% host genes of the aberrantly methylated lncRNAs were also aberrantly methylated. Taken together, these results suggested that the aberrant methylation was associated with ncRNAs in breast cancers. And the genome wide study not only discovered many intergenic ncRNAs that cannot be detected by microarray, but also extended the intragenic ncRNA candidates. Our data provided novel evidence of the mechanisms behind ncRNA dysregulation in breast cancers.

### Aberrant epigenetic regulation patterns of ncRNA promoters in the context of CGIs

Although more and more ncRNAs were found to be epigenetically dysregulated in various types of cancers, the genome-wide aberrant methylation of ncRNAs in breast cancer is poorly revealed. Recently, Weber et al. reported that gene promoters with weak CGIs unbound to RNA polymerase II were frequently methylated (Koga et al. 2009). However, the relevance of specific features of DNA methylation, such as the location of aberrant methylation, or whether with CGIs, to the development of cancers remains poorly understood. Profiling genome-wide DNA methylation of promoters is expected to produce different patterns of promoter aberrant methylation. Above DMR analyses revealed that the ncRNA promoters’ aberrant methylation was involved in breast cancer development, and thus we next looked for the aberrant methylation pattern around transcription start sites (TSSs) of ncRNAs.

Firstly, visualization of the aberrant methylation marks in the context of CGIs revealed the presence of several distinct aberrant methylation patterns on lncRNA promoters. Broadly, these aberrantly methylated lncRNA promoters fell into two main groups based on the presence or absence of a CGI within their promoters. Interestingly, although 34.81% (n=473) of the hypermethylated lncRNA promoters lacked CGIs, they exhibited aberrant hypermethylation around the TSSs (Figure 4A). The remaining 65.19% (n=886) hypermethylated lncRNA promoters had CGIs and four distinct aberrant methylation patterns were observed (Figure 4B and Table S1): (1) aberrant methylation was mostly confined to CGIs; (2) aberrant methylation was positioned to the 5’ shore of CGIs; (3) aberrant methylation was positioned to the 3’ shore of CGIs and (4) aberrant methylation was overlap with CGIs or on both shores of CGIs. We found that aberrant hypermethylation was mostly confined to the CGIs or 5’/3’ shore of the CGIs (Figure S3). Next, we computed the average aberrant methylation frequency around the TSSs and observed that the aberrant hypermethylation frequency was particularly high immediately downstream of the TSSs in these hypermethylated promoters (Figure 4A). This region was generally regarded as core promoter (Bajic et al. 2004), in which RNA polymerase II is recruited to the DNA and acts to initiate transcription. Aberrant methylation in core promoter regions may lead to transcriptional inactivation/activation of cancer-related genes and plays an integral role in tumorigenesis. For the hypomethylated lncRNA promoters in breast cancers (Table S2), we found that nearly 89.91% were lack of CGIs (Figure 4C and Figure S3), the aberrant hypomethylation frequency peak was immediately downstream of TSSs (Figure 4C), which was similar to hypermethylated promoters. However, the aberrant methylation frequency of hypomethylated promoters with CGIs was distinct from other promoters, showing a lowest aberrant hypomethylation frequency around the TSSs (Figure 4C). Next, we investigated whether any subtypes of lncRNAs were preferentially to aberrantly methylated in breast cancers. As a result, intragenic lncRNAs were observed to contribute a large slice in each pattern of aberrantly methylated lncRNA promoters. In addition, the intergenetic lncRNAs were overrepresented in hypomethylated lncRNAs (Figures 4B and 4D).

**Figure 4.**
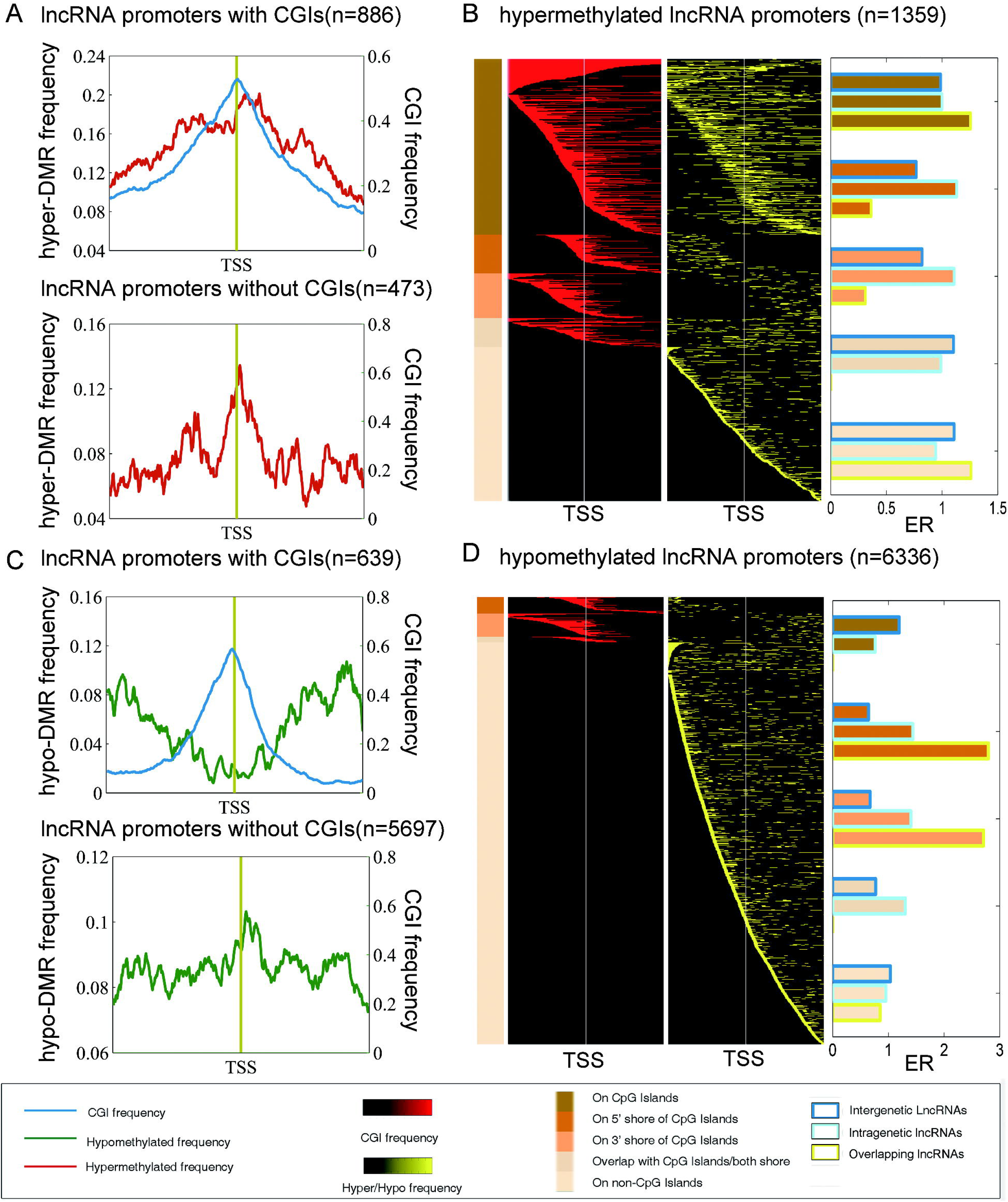
Aberrant methylation patterns around the TSSs of lncRNAs in the breast cancers. (A) Average aberrant hypermethylation frequency of 886 lncRNA promoters with CGIs and 473 lncRNA promoters without CGIs. (B) Heat map of CGI frequency and aberrant hypermethylation frequency in breast cancer. A total of 1359 lncRNA promoters that harboured hypermethylation (yellow) were shown. Each row represents a unique promoter region at 10-bp window size, covering ±2kb flanking the transcription start sites. The location of a CGI (red) in the aberrantly methylated lncRNA promoters is shown in the first column. Promoters are ordered by the location of methylation on a CGI, adjacent to the island (shore) or promoters that lacked CGI as represented with different shades of brown on the left. (C) Average aberrant hypomethylation frequency of 639 lncRNA promoters with CGIs and 5697 lncRNAs that were lacked of CGIs. (D) Heat map of CGI frequency and aberrant hypomethylation frequency in breast cancer. The ratio of lncRNA subtypes for each pattern was shown adjacent the heat map.

Besides lncRNAs, genetic alterations and failure of posttranscriptional regulation might cause the dysregulation of subsets of miRNAs, but epigenetic alterations also appear to be likely culprits (Lujambio et al. 2008). We found that 278 miRNAs were hypermethylated (Table S3) and 802 were hypomethylated in the promoter regions (Table S4). Similar to lncRNAs, aberrant hyper-methylation of miRNA promoters were confined to CGIs while hypo-methylation miRNA promoters were lack of CGIs (Figures S4 and S5). Comparing the aberrant methylation frequency distributions between miRNAs and lncRNAs, we found that the distribution of miRNAs was similar to lncRNAs (Figures S4 and S5), suggesting similar epigenetic mechanism between different kinds of ncRNAs. The aberrantly methylated frequency, especially hypermethylation frequency, was also higher around TSSs, implying the miRNA promoters were frequent targets of aberrant DNA methylation in human breast cancer (Vrba et al. 2013). Among these identified miRNAs, many have been demonstrated to be involved in the process of tumorigenesis, such as *hsa-mir-29b* (Suzuki et al. 2012), *hsa-mir-21* (Yan et al. 2008) and *hsa-mir-10b* (Ma et al. 2007).

### Regulation of ncRNAs expression by DNA methylation

To identify aberrant methylation of ncRNAs with functional significance, we next examined ncRNA promoter methylation events associated with transcriptional changes. Two scenarios of particular biological relevance were considered. In the first scenario, lower expression of ncRNA is due to hypermethylation (silencing), and in the second, higher expression of ncRNA is due to hypomethylation (activating). Considering 13,463 lncRNA genomic loci, 533 hypermethylated lncRNAs were down-regulated in MCF-7 cell line, whereas 829 hypomethylated lncRNAs were up-regulated in MCF-7 cell line. Meanwhile, we observed that 219 hyper- and 1,683 hypomethylated lncRNAs were associated with transcription changes in HCC1954 cell line. As breast cancer is a highly heterogeneous disease and the cell lines used here represent two distinct subtypes. Next, we explored these aberrantly methylated lncRNAs to identify the common ones. Finally, 156 hyper- and 448 hypomethylated lncRNAs candidates were derived in two cell lines (Figure 5A).The identification of this set of lncRNAs in two cell lines suggests that the lncRNAs are actually dysregulated in breast cancers. Among these lncRNAs, we found that they were distributed in various patterns of aberrantly methylated lncRNAs. However, in consistence with previous studies in coding genes, the aberrantly hypermethylated lncRNAs with CGIs were more likely to be associated with transcriptional repression. About 54.45% hypermethylated lncRNAs with CGIs were associated with transcriptional repression in MCF-7 cells and HCC1954 cells (Figure 5A). As both DNA methylation and histone modification are involved in establishing patterns of gene expression during the progression of caners, next we explored whether these aberrant expression of lncRNAs were also associated with altered histone modification. The occupancy of aberrant H3K4me3 and H3K27me3 modifications were assessed in MCF-7 and HCC1954 by comparing with Human Mammary Epithelial Cells (HMEC). A notable observation is that about 50% of the hypermethylated lncRNAs display reduced H3K4me3 or elevated H3K27me3, meanwhile, about 30% of the hypomethylated lncRNAs were with elevated H3K4me3 and reduced H3K27me3 (Figure 5B). However, the aberrant DNA hypermethylation of ncRNAs is likely to co-occur with aberrant H3K4me3 modification than H3K27me3. In the contrast, ncRNA hypomethylation is associated with the alteration of H3K27me3 levels. These are consistent with the previous observations that H3K27me3 and DNA methylation are mutually exclusive, especially for CGIs, and total loss of DNA methylation is associated with alteration of H3K27me3 levels in embryonic stem cells (Brinkman et al. 2012). To visualize the epigenetic patterns at these aberrantly methylated lncRNAs, H3K4me3 intensity plots showed the occupancy in three breast cell lines. We found that the aberrant histone modification is extremely high at the TSS location, reinforcing the notion that aberrant epigenetic modifications were strongly biased toward core promoters (Figure 5C). Altered DNA methylation is ubiquitous in human cancers and specific methylation changes are often correlated with clinical features. In the next step, we exampled whether these lncRNAs can be used as potential biomarkers for early detection. Importantly, these lncRNAs were capable of distinguishing breast cancer samples from healthy controls, which might be aid in the future development of diagnostic biomarkers for breast cancer. The performance of these lncRNAs was evaluated by area under the receiver operation characteristic (ROC) curve (AUC). ROC curves of the hyper- and hypomethylated lncRNAs gave AUC values of 0.944 and 0.983 in the CMS dataset, respectively (Figure 5D).

**Figure 5.**
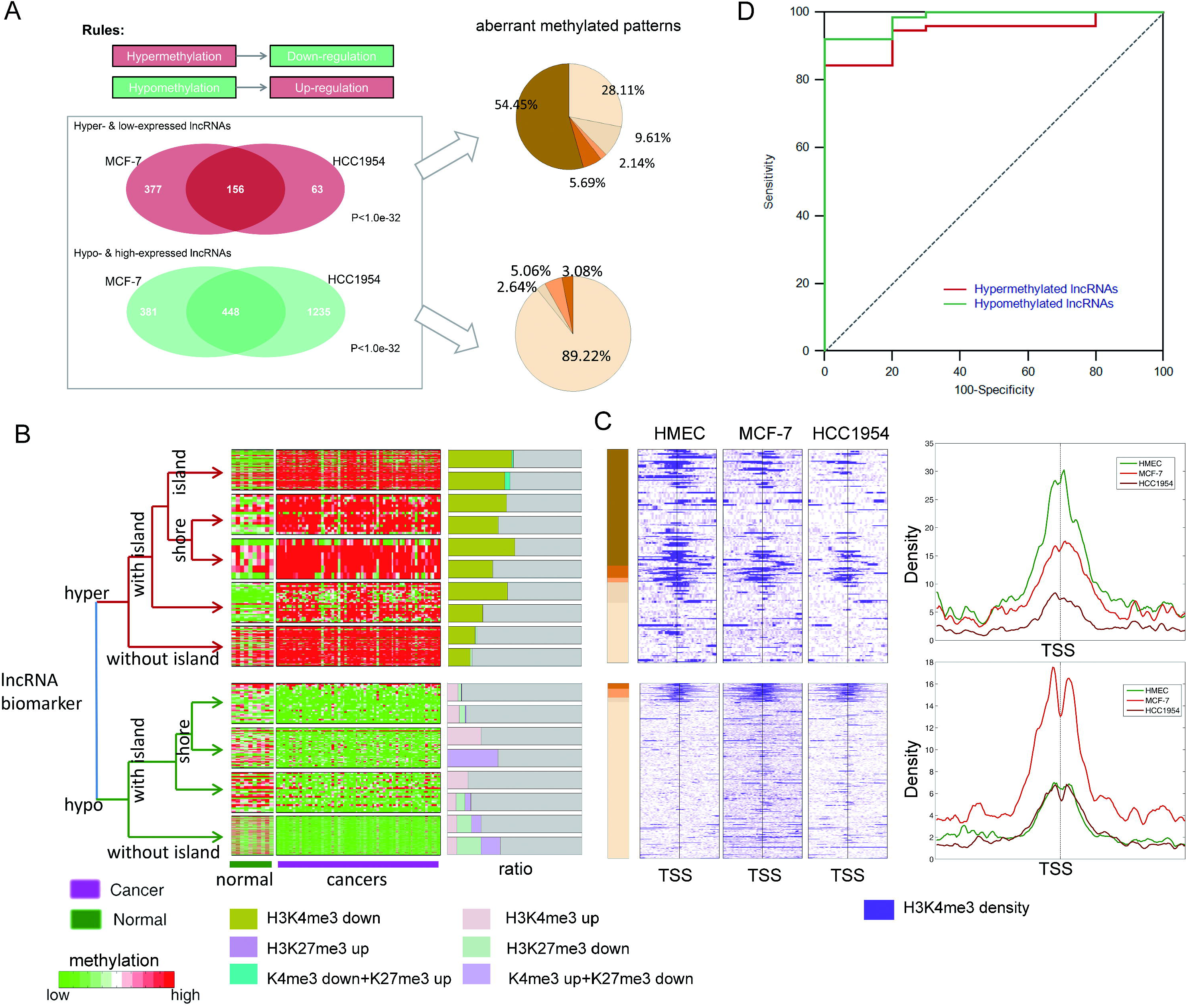
Regulation of lncRNA expression by DNA methylation in breast cancers. (A) The aberrantly methylated and expressed lncRNAs in two cell lines were significantly overlapped. (B) Clustering map of the selected lncRNA biomarkers in CMS methylation. The lncRNAs were grouped based on the aberrant methylation patterns. The barplot adjacent the heat map shows the percentage of lncRNAs in that group covered with aberrant histone modification. (C) Colour profiles of the H3K4me3 sequencing read densities from HMEC, MCF-7 and HCC1954. Each profile shows the 4kb regions surrounding the aberrantly methylated lncRNAs. Average profiles are shown on the right of the colour profiles. (D) ROC map of the selected markers.

Several miRNAs have been reported to be expressed aberrantly in breast cancers and might be used as potential biomarkers for cancer diagnosis. Next, we aimed to examine the miRNA expression profiles in tissue samples of breast cancer to explore the clinical significance in disease diagnosis of the aberrantly methylated miRNAs identified above. We then used the miRNA expression in TCGA dataset to select the miRNA biomarkers using the similar method as lncRNAs. As a result, 26 miRNAs (5 hypermethylated and 21 hypomethylated) were selected as biomarkers based on the concordance between differential methylation and differential expression. Of these five hypermethylated miRNAs, three were confined to CGIs. Although most of the hypomethylated miRNAs were lack of CGIs in their promoters, aberrant methylation was positioned to the shores of CGIs of several miRNA promoters, such as *hsa-mir-106a*, *hsa-mir-1292* and *hsa-mir-3613*. When hierarchical clustering of samples using the methylation and expression of the 26 miRNAs (Figures 6A and 6C), most breast cancers were able to be distinguished from benign adjacent tissues, with overall AUCs larger than 0.90 in the CMS and TCGA datasets, respectively (Figures 6B and 6D). Among the 26miRNAs, most were previously found to be associated with breast cancer (Table S5 and Text S1).

**Figure 6.**
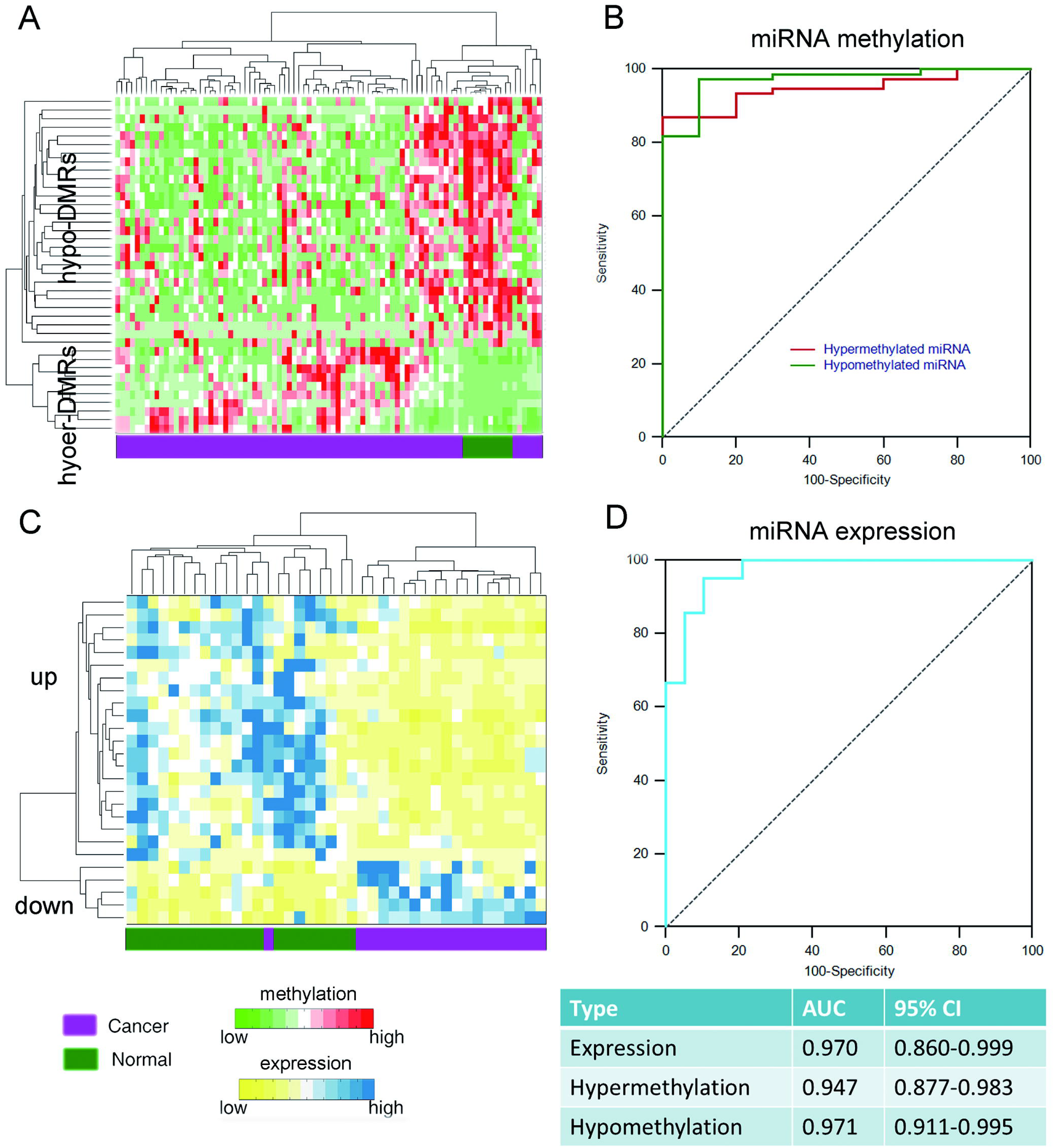
Regulation of miRNA expression by DNA methylation in breast cancers. (A) and (C) Clustering map of the selected miRNA biomarkers in CMS methylation dataset and TCGA expression dataset. (B) and (D) ROC of the miRNA biomarkers.

Together, our findings illustrated how epigenetic mechanisms were linked to aberrant expression of ncRNAs and offered a strong rationale to develop epigenetic-based therapeutics for treatment of breast cancer patients.

### Epigenetically dysregulated ncRNAs disrupt functions associated with breast cancer

Increasing number of lncRNAs are being characterized, however, the functions of most lncRNA genes are still unknown. Besides, functional prediction of lncRNAs is also hampered by the lack of collateral information such as molecular interaction data and expression profiles. The large scale functions annotated to the lncRNAs were mainly predicted by co-expression and genomic adjacency (Liao et al. 2011; Guo et al. 2013). However, despite similar expression patterns, groups of functionally related genes can be further distinguished at the chromatin level (Wamstad et al. 2012). We expected that genes with similar aberrant methylation patterns may have similar functions. To test this idea, we first investigated the aberrant methylation patterns of protein coding genes. As a result, we found that protein-coding genes share similar aberrant methylation patterns with ncRNAs (Details in Text S1 and Figure S6 and S7). Next, we determined gene set functional similarity between genes with distinct aberrant methylation patterns (Figure 7A and Text S1). Consistent with our hypothesis, we found that genes with similar aberrant methylation patterns were with higher function similarity than genes with distinct patterns (Figure 7B). In addition, genes that have the methylation change in the same direction were with higher similarities than those in the opposite direction (Figure S8). As the general large-scale functional annotation of lncRNAs has been based on the ‘guilt-by-association’ principle, we combined this principle and aberrant methylation patterns to predict the probable functions of the aberrantly methylated and expressed lncRNAs identified above. For example, the lncRNA *ENSG00000232821* is hypermethylated confined to CGI. We found that a coding gene-*TWIST1*, adjacent to this lncRNA is also hypermethylated confined to CGI (Figure 7C). *TWIST1* is an important transcription factor that has been implicated in cell lineage determination and differentiation. *TWIST1* promoter methylation has been found significantly more prevalent in malignant compared with healthy breast tissues. It is reasonable to infer that this lncRNA may be also involved in development of breast cancer. We found that the potential functions of this lncRNA in NONCODE were regulation of growth and development, suggesting that this lncRNA played important roles in the development of cancers. Our methylation analysis indicated that *ENSG00000232821* was hypermethylated in breast cancers, providing the epigenetic view to explain its aberrant expression. Next, we aimed to explore the dysregulated pathways of different patterns of aberrantly methylated lncRNA biomarkers. As a result, we found that only the hypermethylated lncRNAs confined to CGIs and hypomethylated lncRNAs lack of CGIs enriched pathways directly associated with the development and progression of breast cancers (Figure 7D). The hypermethylated lncRNAs mainly affected the cell cycle and signaling pathways while the hypomethylated lncRNAs disturbed most of metabolism pathways. Cancer cells require metabolism to sustain their existence, including support of several functions, such as cell maintenance, proliferation and motility. Common to all this activities is the demand for energy (Dolfi et al. 2013). These results indicated that lncRNA hypomethylation may be linked to carcinogenesis by causing the signaling and metabolism pathways dysregulation.

**Figure 7.**
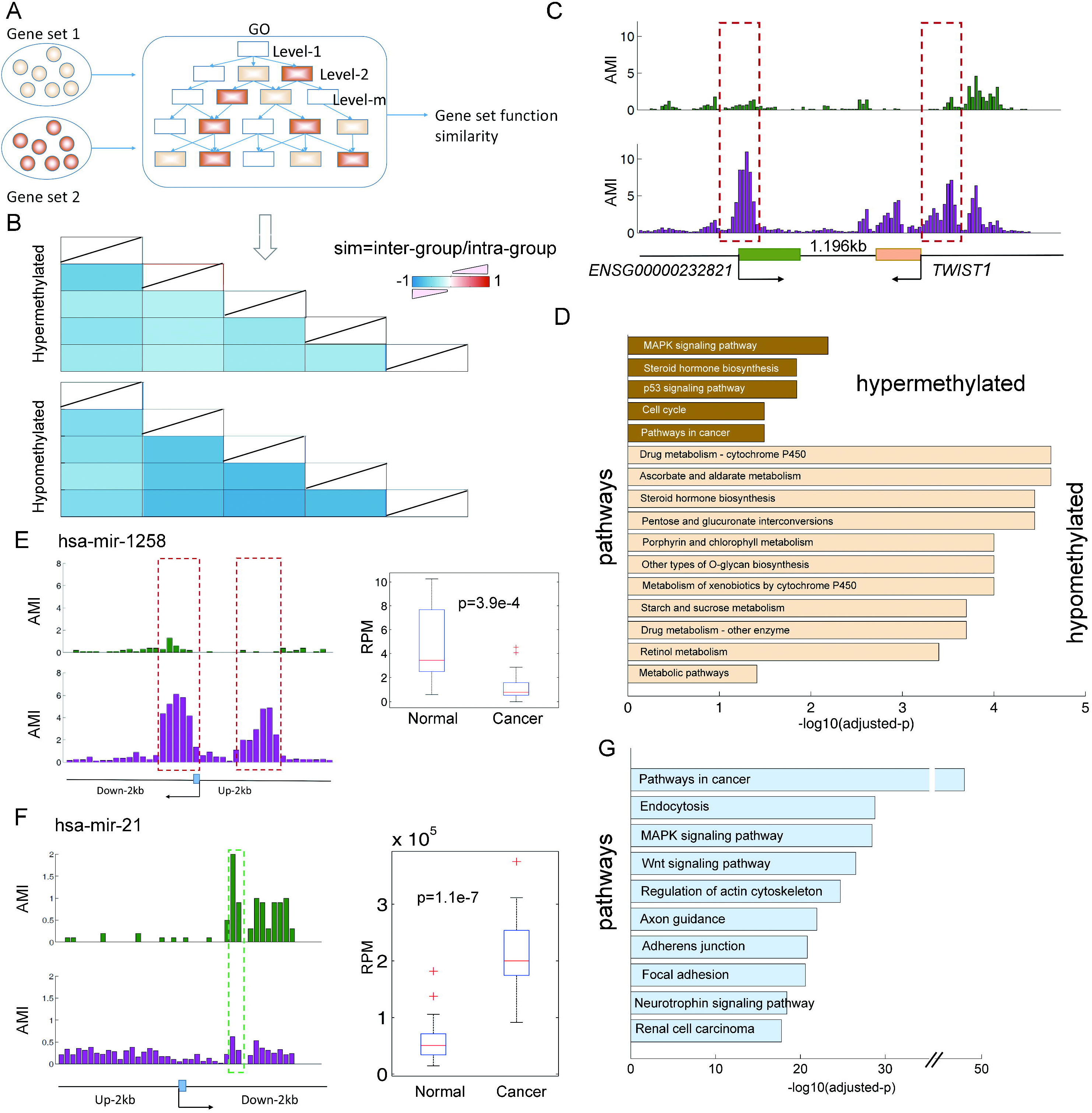
Aberrantly methylated ncRNAs disturb widely functions associated with breast cancer. (A) A pipeline to compute the gene set functional similarity. (B) Genes with similar aberrant methylation patterns were with high functional similarity. The similarity score was normalized to the intra-classes. (C) An example of the lncRNA and protein-coding gene. The average methylation intensities (AMI) in the normal and breast cancer samples were shown. Green, normal samples; Pink, cancer samples. (D) The biological processes enriched by adjacent genes of lncRNA with similar aberrant methylation patterns. (E) The average methylation and expression levels of *hsa-mir-1258* in the normal and breast cancer samples. RPM, reads per million. (F) The average methylation and expression levels of *hsa-mir-21* in the normal and breast cancer samples. (G)The top ten pathways enriched by targets of miRNA biomarkers.

MiRNAs play their important roles through targets; functional enrichment analysis revealed that the predicted targets of the 26 aberrantly methylated and expressed miRNAs, greater than that expected by chance, were implicated in functions which have important roles in direct tumor growth and metastasis (Figure 7G, Table S6). *hsa-miR-29b* was observed to be highly over-expressed in breast cancer, and enhanced *hsa-miR-29b* impaired apoptosis, increased tumor cell migration and invasion directly by targeting *PTEN* tumor suppressor (Wang et al. 2011). In our present study, we found that although the promoter of *miR-29b* lack of CGI, it was hypomethylated in breast cancer (Table S4), providing the epigenetic view of aberrant expression of *hsa-miR-29b* in cancers. Another example is the tumor suppressor-*hsa-miR-1258*, which was hypermethylated on 3’ shore and lowly expressed in our current study (Figure 7E). Heparanase (HPSE) is a potent protumorigenic, proangiogenic, and prometastatic enzyme that is over-expressed in brain metastatic breast cancer (BMBC). Evidences have shown that *hsa-miR-1258* inhibited the expression and activity of heparanase in BMBC cells, stable expression of *hsa-miR-1258* in BMBC cell inhibited heparanase in vitro cell invasion and experimental brain metastasis (Zhang et al. 2011a). In addition, high *hsa-miR-21* expression was associated with mastectomy, larger tumor size, higher stage, higher grade, estrogen receptor status. Increasing studies have identified many downstream targets of *hsa-miR-21* in various types of cancers, such as *WNT5A*, *PTEN* and *BRCA1* (Yan et al. 2008). Consistent with previous studies, the *hsa-miR-21* is significantly highly expressed in breast cancers in our analysis (p=1.1e-7). However, the mechanism underlying *hsa-miR-21* high expression remains largely unknown. We found that the promoter of this miRNA lacks of CGI and is hypomethylated in breast cancer (Figure 7F), suggesting a possible epigenetic role in regulating its activity in breast cancer. Moreover, we found that some of the targets of this miRNA are hypermethylated, such as *BRCA1*, *CDK6* and *WNT5A*. Mutation of the *BRCA1* tumor suppressor gene is an important contributing factor in hereditary breast cancer; however, *BRCA1* mutations have not been detected in some types of breast cancers, suggesting that DNA methylation and/or miRNA repression of *BRCA1* may participate in the genesis of breast cancers (Rice et al. 2000). Our analysis indicated that hypomethylation of the miRNA regulators and hypermethylation of the gene may be another repression mechanism of this tumor suppressor gene.

Taken collectively, these observations suggest that the identified ncRNAs in our study play a critical role in the development and progression of breast cancer, systematical investigation of their functions may provide new potential targets for breast cancer therapy.

### Noncoding-RNAs and coding genes coordinately mediated pathways dysregulation in breast cancers

Tumorigenesis is a complex dynamic biological process that includes multiple steps of genetic and epigenetic alterations, aberrant expression of ncRNA, and changes in the expression profiles of coding genes (Cao et al. 2013). In addition, ncRNAs are integral components of biological networks with fundamental roles in regulating gene expression. However, our understanding of the complex network intertwined by coding genes and ncRNAs in cancer biology is still in an early stage. Our above analyses have revealed that the aberrant methylation of ncRNAs in breast cancer could perturb various pathways. In addition, we found that the aberrant methylation of lncRNA or miRNA can both perturb some common pathways, such as cell cycle, MAPK signaling pathways, indicating that ncRNAs mediate the dysregulation of pathways in a coordinate manner. As one of the most important cellular processes, the cell cycle is under precise regulation in all organisms. Mis-regulation of the cell cycle can lead to catastrophic cellular events, e.g. premature apoptosis or abnormal proliferation of cells, which are the causes of cancers (Cheng et al. 2013). We found that both the lncRNA and miRNA regulated the key components of this pathway. One example is the gene *CDK6*, which is responsible for modulation of the activities of *Rb* family growth-suppressing proteins. Evidences have shown that normal human mammary epithelial cells have a high amount of *CDK6* activity, but all breast tumor-derived cell lines had reduced levels, with several having little or no *CDK6* (Lucas et al. 2004). Our analyses demonstrated that DNA methylation mainly be one mechanism for repression of this cell cycle associated genes (Figure 8A). Moreover, we found that this gene was also targeted by two hypomethylated miRNAs, *hsa-miR-21* and *hsa-miR-29b*. The hypomethylation of these two key regulators may further repress the activity of *CDK6*, implying the complement effect of DNA methylation and miRNA regulation. Another example is the hypermethylated gene-*CCND2* (Figure 8A), which was also regulated by a hypomethylated miRNA (*hsa-miR-16*). This cyclin forms a complex with and functions as a regulatory subunit of *CDK4* or *CDK6*, whose activity is required for cell cycle G1/S transition (Pic-Taylor et al. 2004). In addition, two hypermethylated lncRNAs share the similar aberrant methylation pattern with this gene (both hypermethylated on CGIs), implying that these two lncRNAs play key roles in cell cycle regulation.

**Figure 8.**
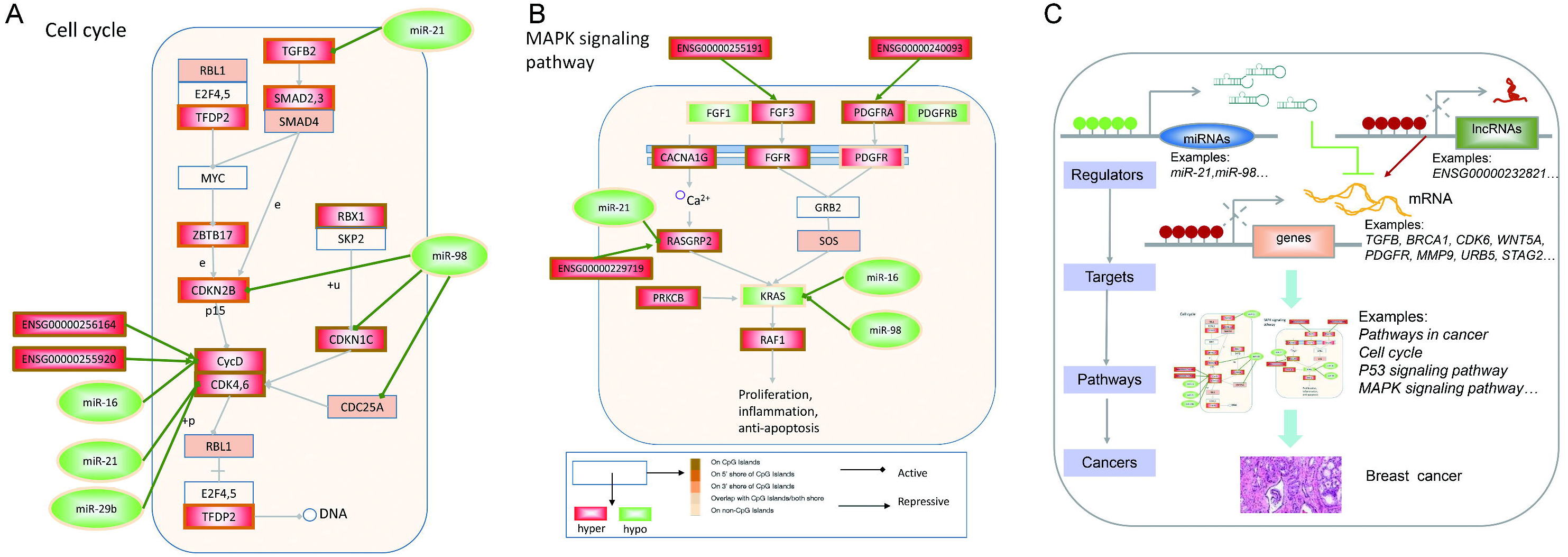
NcRNAs and coding genes coordinately mediated the pathway dysregulation in breast cancers. (A) The cell cycle pathway dysregulated by ncRNAs and coding genes. Hypermthylated genes were marked in red and hypomethylated genes were marked in green. The aberrantly methylated patterns were marked on the edge of the rectangle. The experimentally validated miRNA regulations from TarBase and adjacent lncRNAs with common aberrant methylation patterns were linked. (B) MAPK signalling pathway. (C) A model illustrating the dysregulated cellular network intertwined by ncRNAs and coding genes, ncRNA and coding genes coordinately mediated the pathway mis-regulation.

MAPK signaling pathway is evolutionarily conserved kinase modules that link extracellular signals to the machinery that controls fundamental cellular processes such as growth, proliferation, differentiation, migration and apoptosis (Dhillon et al. 2007). It is revealed in our study that most of genes in this pathway were aberrantly methylated in breast cancer (Figure 8B). Among these genes, some were reported to be involved in tumorgenesis, such as *FGFR1*, *KRAS* and *PRKCB*. Although there are correlations between the expression of FGFRs and breast cancer progression (Hynes 2000), the underlying mechanism of its over-repression is still not fully understood and may, in fact, be multiple. Our study indicated that DNA hypomethylation may be another mechanism response for its over-repression in breast cancers. The Ras oncogene family has been very extensively studied and the fundamental implication of Ras protein in pathological processes such as cancer and development (Fernandez-Medarde and Santos 2011). The experimental observations of accumulated for many years document that somatic mutations are the typical genetic lesions affecting Ras. However, the mutational event for *RAS* gene does not seem to be involved in some breast cancer patients (Sanchez-Munoz et al. 2010). We found that the *KRAS* gene is under strict regulation by DNA methylation and miRNAs. Besides hypomethylation of *KRAS* gene itself, two miRNAs (*hsa-miR-16* and *hsa-miR-98*) that regulated *KRAS* were also found to be aberrantly methylated. Dysregulation of these two miRNAs have also been implicated in the development of breast cancer (Cho 2007). These observations implied that contrary to mutation of *KRAS*, the epigenetic dysregulation of ncRNA regulators and itself may be another mechanisms response for aberrant expression.

Cells employ cellular signaling pathways and networks that are intertwined by non-coding and coding RNAs to drive biological processes. Our observations indicated that aberrant methylation of ncRNA regulators and target mRNAs, in principle, might result in aberrant expression of genes, which may encourage neoplastic transformation (Figure 8C). Epigenetic dysregulation of ncRNAs and mRNAs coordinately mediated the pathways mis-regulation, which then leads to the development and progression of breast cancer. Together, integration of multi-dimensional genomic data is likely to generate more insight into the fundamental pathway wiring in cancer, with the more focused aim of identifying candidate ncRNAs and coding-genes induced by epigenetic alterations that will be useful for interpreting complex mechanisms underlying the cancer genome.

## DISCUSSION

In this study, we characterized genome-wide methylation patterns in breast cancer using a novel high throughput sequencing methodology. Our study reveals important DMRs and aberrant methylation patterns of ncRNAs in breast cancers. Consistent with prior studies in breast cancer cell lines (Ruike et al. 2010), we found widespread hypomethylation and local hypermethylation in breast cancers. Besides the extensive CGI hypermethylation, an increasing number of ncRNA promoters were found to be aberrantly methylated, including lncRNAs and miRNAs. Among these ncRNAs, intergenic lncRNAs were observed to contribute a large slice of the aberrantly methylated lncRNA promoters in breast cancers. Our comparative methylome analyses allow us to dissect the genome-wide methylation patterns of ncRNAs. We summarized five patterns of ncRNA promoter aberrant methylation, where aberrant methylation not only span the CGI, but also in 5’ and 3’ shores, which have been found previously in colorectal cancer (Irizarry et al. 2009) and prostate cancer (Kim et al. 2011) for protein coding genes. Although coding and non-coding RNAs function in different ways, they share similar aberrant methylation patterns.

Integrating epigenomics data with expression profiling of ncRNA, we found that the aberrant methylation of ncRNAs were associated with transcriptional changes. Numerous studies show that it seems obvious that the expression of ncRNAs can also be modified by genetic variants, such as copy number variants (CNV) and single nucleotide polymorphism (SNP) (Ramsingh et al. 2013). However, we found that about 11.8% and 9.71% of hypermethylated lncRNAs and miRNAs were located in the regions of recurring deletions (Beroukhim et al. 2010). This result is consistent with recent observations that although individual genetic alterations appear to affect the gene expression and methylation levels, these are rare (Gibbs et al. 2010). These observations demonstrated that ncRNA expression can be controlled not only by genetic but also epigenetic determinants, which is consistent with recent observations (Aure et al. 2013; Jacobsen et al. 2013). Next, we identified a panel of ncRNA biomarkers that can effectively discriminate cancer and normal samples. Finally, functional analysis indicated that the aberrant methylation of ncRNAs disturbs widely processes associated with the development and progression of breast cancers.

Polycomb targeted genes are frequent targets of aberrant DNA methylation in cancers, and some miRNA genes have also been showed to be targeted by polycomb proteins. Therefore, we examined whether polycomb targeted ncRNAs are more likely to be targets of aberrant DNA methylation in breast cancer. To this end, we used the repressed regions of Ernst et al. study as polycomb targeted regions (Ernst et al. 2011). We found a significant overlap between ncRNA promoters occupied by the polycomb and those aberrantly hypermethylated in cancer cells (Figure S9). The comparisons show that ncRNA promoters targeted by polycomb were more likely to be hypermethylated in cancer. These observations provided additional evidences for the link between polycomb repression and aberrant DNA methylation in breast cancer. The high portion of polycomb targets among ncRNA promoters may, to a certain extent, explain the high proportion of ncRNA promoters with aberrant methylation in cancer. However, it is possible that no direct functional link between both phenomena. Such ncRNAs are unmethylated in normal cells and repressed by polycomb proteins, and acquire DNA methylation as an alternative silencing mechanism. These ncRNAs, have tumour suppressive functions and are inducible in normal cells, although repressed by polycomb. One can hypothesize that the epigenetic switch to DNA methylation-mediated repression reduces the plasticity of the regulatory program of these ncRNAs, locking silencing of the key ncRNA regulators and contributing to the cell’s abnormal growth potential.

Emerging evidence suggests that the vast, often uncharted, noncoding landscape of human genome plays an important role in biology and cancer progression. Despite growing appreciation of the importance of ncRNAs in breast cancer, our knowledge of the functions of ncRNAs remains limited. MiRNAs have well-established differential patterns of expression in cancer and function as tumor suppressors or oncogenes by silencing target gene expression. Much less is known about lncRNAs, subsets of which have been characterized as epigenetic factors, enhancers and antisense transcripts. However, the functions of most lncRNAs in cancer remain a mystery. Currently, the large scale functions annotated to the lncRNAs mainly predicted by co-expression and genomic adjacency. In addition, we found that genes with similar aberrant methylation patterns tend to have higher functional similarity and then the functions of aberrantly methylated lncRNAs were predicted based on similar patterns. Our results have shown that groups of functionally related genes can be further distinguished at the chromatin level, providing an epigenetic view to dissect the functional roles of lncRNAs in complex diseases. Although coding and noncoding genes function in different manners, our analysis highlighted at least two cases that ncRNAs and coding genes coordinately mediated the pathway dysregulation in the development of breast cancers. Systematically analysis of the combinations in the context of pathway is likely to generate more insight into the fundamental network wiring in cancer.

In contrast to hypermethylation, which seems to happen locally, preferentially targeting the promoters of genes, the mechanism of global hypomethylation is a long-standing question in cancer epigenetics (Hon et al. 2012). Our results seem to support that hypomethylation appears to target large inter-genic repeat regions. To assess the frequency of hypomethylation at repeats, we examined the ratio of repetitive elements covered with DMRs. Dramatically, more repetitive elements were covered by hypo-DMRs than hyper-DMRs (Figure S10A). Additionally, the hypo-DMRs were significantly over-represented in SINEs and LTRs than random regions (p<0.001, Figure S2D). Increased expression of repetitive elements is a hallmark of cancer cells, and our current and previous studies suggest a possible mechanism that this aspect of cancer may be attributed to global hypomethylation (Hon et al. 2012). Analysis of GO and KEGG functional categories for the adjacent genes of these repetitive elements revealed overrepresentation of functional categories common in cancer development and progression, such as cell adhesion molecules, MAPK signaling pathway and VEGF signaling pathway (Figure S10B-H). A well-recognized result of DNA hypomethylation of these repeat elements are the induction of global genetic instability and, though the evidence is mostly indirect, this has been observed in cases of extreme global hypomethylation in murine models and in multiple studies of human cancers (Rodriguez et al. 2006). We supposed that DNA hypomethylation-related instability may disturb the functions of adjacent genes, thus leading to the dysregulation of pathways.

In summary, we have performed a comparative methylome analysis of breast cancer and normal controls and provided the precise genomic locations that undergo methylation changes. Our data shows the suitability of MBDCap-seq to investigate cancer methylomes and to identify novel epigenetically dysregulated ncRNAs (lncRNAs or miRNAs). This study presents the aberrant methylation patterns of ncRNAs, which will be a highly valuable resource for investigations at understanding epigenetic regulation of breast cancers. In addition, the identified ncRNA biomarkers in our study might be potentially used for diagnostic purposes. Together, these results demonstrate the ability of integration of multi-dimensional epigenomic and transcriptomic data to identify candidate ncRNAs induced by epigenomic alterations, which will be useful for interpreting complex cancer genome.

## MATERIAL AND METHODS

### Genome-wide DNA methylation profiles

The genome-wide DNA methylation profiles from the Cancer Methylome System (CMS, http://cbbiweb.uthscsa.edu/KMethylomes) were used in the current study (Gu et al. 2013). The CMS consists of whole genome-wide methylation data conducted by the methyl-CpG binding domain proteins followed by sequencing, or MBDCap-seq protocol. The current version contains methylation profiles from endometrial and breast cancers, and some normal samples are included for comparison purpose. We used the methylation intensity profiles for 77 breast cancer and 10 normal samples. Briefly, sequencing reads in 36bp lengths were mapped by the ELAND algorithm, with up to two mismatches. Genome-wide methylation intensity at 100 base-pair resolution was quantified by the read numbers that were located within the genomic bin, and then normalized based on the unique read numbers for each sample by the linear method (Derrien et al. 2012). The following equation was used for linear normalization:

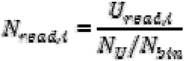

Where *N_read,i_* is the normalized read number of the genomic bin *i*, and *U_read,i_* is the unique mapped read number of genomic bin *i*, *N_U_* is the total unique mapped reads for the sample, and *N_bin_* is the total bin number of human genome.

### Genome-wide expression of ncRNAs

As the expression profiles of lncRNA in breast cancer tissues are insufficient currently, we used two breast cancer cell lines (MCF-7 and HCC1954) and one control (HMEC) to analyze the expression. The raw reads data of MCF-7 and HMEC were downloaded from the ENCODE project (Bernstein et al. 2012) and the data for HCC1954 was kindly provided by Hon et al (Hon et al. 2012). To quantify the FPKM expression (reads per kilobase of exon model per million mapped reads) at lncRNAs, the Cuffdiff program was applied to the mapped TopHat reads (Trapnell et al. 2010). Only the subset of RNA-seq reads that are uniquely mapped to hg18 were used.

In order to investigate the effects of methylation on expression, miRNA expression analysis was performed using publically available dataset. Twenty-one tumor-adjacent normal samples from The Cancer Genome Atlas (TCGA) were used. These pairs were with both DNA methylation and gene expression profiles (Table S7). The gene expressions were evaluated with the AgilentG450 microarray, and we directly downloaded the log transformed data. A total of 17,814 genes are represented on this array. In addition, among the twenty-one pairs of samples, twenty breast cancer and nineteen normal samples with miRNA expressions were used in the miRNA biomarker analyses. The miRNA expression were measured by miRNA-Seq. Mapping the reads to the human genome, the reads per million mapped to each miRNA were used to represent the expression of miRNAs. As a result, a total of 1046 pre-miRNAs were analyzed.

### Annotation of genomic features

Long noncoding RNAs (lncRNAs) have been shown to play important roles in many biological processes, including cancers. The GENCODE annotation represents a valuable resource for studies of lncRNAs. We downloaded the human lncRNA annotation from GENCODE (version 18), including 13,479 gene loci. After converting the genomic location to hg18 by the LiftOver in UCSC, 13,459 lncRNAs were used for analyses. In addition, these lncRNAs were further reclassified into three biotypes based on their location with respect to protein-coding genes: intergenic lncRNA, intragenic lncRNA and overlapping lncRNA. The promoters of lncRNAs were also defined as 2kb on either side of the TSSs. About 1866 pri-miRNA genomic coordinates were taken from miRBase database (version 20.0) (Griffiths-Jones et al. 2006). Currently, only a small number of human miRNA genes have confirmed transcription start sites (TSSs), however, numbers of studies have shown that miRNA shares common transcriptional patterns with protein coding gene (Ozsolak et al. 2008). In the current study, miRNA promoters were defined similar as protein coding genes, which were defined as 2kb upstream of and 2kb downstream from the start sites of pri-miRNAs.

Coordinates of genomic features, such as refseq genes, intron and exon, are mainly taken from the UCSC database. Similar as previous studies, the gene promoters were defined as 2kb upstream of and 2kb downstream from the TSS. In addition, the genomic coordinates of human CpG islands (CGIs) were also downloaded from the UCSC (Gardiner-Garden and Frommer 1987). CGIs were predicted by searching the sequence one base at a time, and each segment was evaluated for the following criteria: (1) CG content of 50% or greater; (2) length greater than 200bp; (3) CpG ratio greater than 0.6. The 5’-and 3’-shores of CGI were defined as 2kb on either side of the CGI according to previous studies (Irizarry et al. 2009). The list of repeat elements predicted by the RepeatMasker program was also downloaded from the UCSC genome browser and then classified into different groups according to their annotation in the UCSC.

### DMR identification

To detect regions of differential methylation between breast cancer phenotypes and non-neoplastic controls, we screened genomic bins of 100bp. For each bin, the normalized methylation intensity of this bin was compared between sample cohorts with Wilcox rank-sum test. Bins with adjusted p-value less than 0.01 and fold differences greater than twice were identified as differentially methylated bins (Benjamini 1995). For each differentially methylated bin, we defined hyper-methylation if the average normalized methylation intensity of breast cancer group is higher than that of controls, and vice versa (hypo-methylation). Consecutive hyper/hypo-bins with no gap between them were merged into differentially methylated regions (DMRs).

To investigate whether certain structural genomic features were enriched DMRs, we firstly computed the ratios for any given structural genomic features that covered with DMRs. A DMR was regarded as located with a given genomic features if 50% of the DMR was overlapped with the feature. Significance of this ratio was evaluated using random features. The same numbers of random features were generated with the BEDTools by randomly permuting the locations of features within a genome (Quinlan and Hall 2010). The significant p-value is the fraction of the features covered with DMRs in random conditions, which is greater than the value in the real condition.

### Identification of the patterns of aberrant methylation

To investigate the aberrant methylation patterns around the TSSs, each promoter region was firstly divided into 10bp windows. And then the aberrant methylation frequency was calculated for each window, which was defined by the number of base-pairs covered by DMRs divided by the length of window. Similarly, CGI frequency for each window was calculated. Specifically, we focused on two types of aberrantly methylated promoters: aberrantly hyper-methylated and hypo-methylated in breast cancer. Then their aberrant methylation patterns were analysed in the context of CGIs. Finally, five methylation patterns were identified for promoters aberrantly methylated in breast cancer. (1) The aberrant methylation was mostly confined to the CGIs, that was the number of overlapped windows between aberrantly methylated and covered by CGIs was larger than either half of the number of aberrantly methylated windows or that of covered by CGIs; (2) The aberrant methylation was positioned 5’ to the CGIs; (3) The aberrant methylation was positioned 3’ to the CGIs; (4) The aberrant methylation was overlapped with CGIs. (5) The aberrantly methylated promoters lack of CGIs. In these analyses, all the hyper- and hypo-DMRs were analyzed, respectively. The average aberrant methylation frequency or CGI frequency of all the aberrantly methylated promoters was used to obtain the aberrant methylation patterns around the TSS.

### Regulation of ncRNA expression by DNA methylation

A two-sample t-test was carried out to select the differentially expressed miRNAs. The P values were adjusted by the Benjamini and Hochberg (BH) correction procedure to account for multiple tests with the false discovery rate (FDR) < 5%. In addition, the differentially expressed lncRNAs were identified by the fold change method, lncRNAs with fold change greater than 2 or less than 0.5 were identified as over-repressed or under-expressed, respectively.

### Histone modification of ncRNAs in breast cancers

ChIP-seq data (H3K4me3 and H3K27me3) for HMEC and MCF-7 was downloaded from ENCODE and the ChIP-seq data for HCC1954 cell lines were obtained from NCBI Gene Expression Omnibus (GEO) database (GSE29069) (Hon et al. 2012). We downloaded the fastq format data and then the reads were mapped to human genome (hg18) by the software Bowtie (Langmead and Salzberg 2012), and only reads that at most three mismatches were retained for subsequential analysis. The ChIPDiff program (Xu et al. 2008) was used for quantitative comparison of the histone modification levels in the breast cancer cells and the normal cell lines. It employs a hidden Markov model (HMM) to infer the states of histone modification changes at each genomic location. The genomic 1kb bins were first identified as differential histone modification sites (DHMS) and consecutive DHMSs with no gap between them were merged into DHMS regions. And then lncRNAs covered by DHMS regions were regarded as histone modification dysregulation.

### Classification algorithm and validation of ncRNA biomarkers

Integration of methylation and expression, we identified ncRNA biomarkers. In order to evaluate these biomarkers, first, we performed Z-score transformation on the methylation or expression levels across the samples for each of the miRNA and then summarized the Z-scores for these miRNAs in the signature. And then the receiver operation characteristic (ROC) curve is used for classifier evaluation which is drawn by plotting sensitivity against the false-positive rate. The area under the ROC curve (AUC) can be used as a reliable measure of classifier performance. This procedure was performed by the MedCalc program.

### Function enrichment analysis

KEGG pathways and gene ontology (GO) analysis were performed to find enriched pathways using the WebGestalt (Zhang et al. 2005). P-values were multiple tests corrected in order to reduce false-positive rates. Pathways or GO terms with adjusted p-values of <0.01 and with at least two interesting genes were considered significant.

## ACKNOWLEDGMENTS

This work was supported in part by the National High Technology Research and Development Program of China [863 Program, Grant Nos. SS2014AA021102], the National Program on Key Basic Research Project [973 Program, Grant Nos. 2014CB910504], the National Natural Science Foundation of China [Grant Nos. 91129710, 61170154 and 61203264] and China Postdoctoral Science Foundation [2012M520764].

## COMPETING FINANGIAL INTERESTS

The authors declare that there are no potential conflicts of interest were disclosed.

## REFERENCES

Assel S. Issabekova OAB, Vladimir A. Khailenko, Shara A. Atambayeva, Mireille Regnier, Anatoly T. Ivachshenko. 2011. Characteristics of Intronic and Intergenic Human miRNAs and Features of their Interaction with mRNA. World Academy of Science, Engineering and Technology 59.

Augoff K, McCue B, Plow EF, Sossey-Alaoui K. 2012. miR-31 and its host gene lncRNA LOC554202 are regulated by promoter hypermethylation in triple-negative breast cancer. Molecular cancer 11: 5.

Aure MR, Leivonen SK, Fleischer T, Zhu Q, Overgaard J, Alsner J, Tramm T, Louhimo R, Alnaes GI, Perala M et al. 2013. Individual and combined effects of DNA methylation and copy number alterations on miRNA expression in breast tumors. Genome Biol 14(11): R126.

Baer C, Claus R, Plass C. 2013. Genome-wide epigenetic regulation of miRNAs in cancer. Cancer research 73(2): 473–477.

Bajic VB, Choudhary V, Hock CK. 2004. Content analysis of the core promoter region of human genes. In Silico Biol 4(2): 109–125.

Benjamini Y, and Hochberg Y. 1995. Controlling the false discovery rate: a practical and powerful approach to multiple testing. Statistical Society Series B 57: 289–300.

Bernstein BE, Birney E, Dunham I, Green ED, Gunter C, Snyder M. 2012. An integrated encyclopedia of DNA elements in the human genome. Nature 489(7414): 57–74.

Beroukhim R, Mermel CH, Porter D, Wei G, Raychaudhuri S, Donovan J, Barretina J, Boehm JS, Dobson J, Urashima M et al. 2010. The landscape of somatic copy-number alteration across human cancers. Nature 463(7283): 899–905.

Brinkman AB, Gu H, Bartels SJ, Zhang Y, Matarese F, Simmer F, Marks H, Bock C, Gnirke A, Meissner A et al. 2012. Sequential ChIP-bisulfite sequencing enables direct genome-scale investigation of chromatin and DNA methylation cross-talk. Genome Res 22(6): 1128–1138.

Cabili MN, Trapnell C, Goff L, Koziol M, Tazon-Vega B, Regev A, Rinn JL. 2011. Integrative annotation of human large intergenic noncoding RNAs reveals global properties and specific subclasses. Genes & development 25(18): 1915–1927.

Cao W, Wu W, Shi F, Chen X, Wu L, Yang K, Tian F, Zhu M, Chen G, Wang W et al. 2013. Integrated analysis of long noncoding RNA and coding RNA expression in esophageal squamous cell carcinoma. Int J Genomics 2013: 480534.

Cheng C, Ung M, Grant GD, Whitfield ML. 2013. Transcription factor binding profiles reveal cyclic expression of human protein-coding genes and non-coding RNAs. PLoS Comput Biol 9(7): e1003132.

Cho WC. 2007. OncomiRs: the discovery and progress of microRNAs in cancers. Mol Cancer 6: 60.

Croce CM. 2009. Causes and consequences of microRNA dysregulation in cancer. Nat Rev Genet 10(10): 704–714.

Cui P, Lin Q, Ding F, Xin C, Gong W, Zhang L, Geng J, Zhang B, Yu X, Yang J et al. 2010. A comparison between ribo-minus RNA-sequencing and polyA-selected RNA-sequencing. Genomics 96(5): 259–265.

Dai W, Zeller C, Masrour N, Siddiqui N, Paul J, Brown R. 2013. Promoter CpG Island Methylation of Genes in Key Cancer Pathways Associates with Clinical Outcome in High-Grade Serous Ovarian Cancer. Clinical cancer research : an official journal of the American Association for Cancer Research.

Derrien T, Johnson R, Bussotti G, Tanzer A, Djebali S, Tilgner H, Guernec G, Martin D, Merkel A, Knowles DG et al. 2012. The GENCODE v7 catalog of human long noncoding RNAs: analysis of their gene structure, evolution, and expression. Genome Res 22(9): 1775–1789.

Dhillon AS, Hagan S, Rath O, Kolch W. 2007. MAP kinase signalling pathways in cancer. Oncogene 26(22): 3279–3290.

Dolfi SC, Chan LL, Qiu J, Tedeschi PM, Bertino JR, Hirshfield KM, Oltvai ZN, Vazquez A. 2013. The metabolic demands of cancer cells are coupled to their size and protein synthesis rates. Cancer Metab 1(1): 20.

Ernst J, Kheradpour P, Mikkelsen TS, Shoresh N, Ward LD, Epstein CB, Zhang X, Wang L, Issner R, Coyne M et al. 2011. Mapping and analysis of chromatin state dynamics in nine human cell types. Nature 473(7345): 43–49.

Fackler MJ, Umbricht CB, Williams D, Argani P, Cruz LA, Merino VF, Teo WW, Zhang Z, Huang P, Visvananthan K et al. 2011. Genome-wide methylation analysis identifies genes specific to breast cancer hormone receptor status and risk of recurrence. Cancer research 71(19): 6195–6207.

Fernandez-Medarde A, Santos E. 2011. Ras in cancer and developmental diseases. Genes Cancer 2(3): 344–358.

Flintoft L. 2013. Non-coding RNA: Structure and function for lncRNAs. Nat Rev Genet 14(9): 598.

Gardiner-Garden M, Frommer M. 1987. CpG islands in vertebrate genomes. J Mol Biol 196(2): 261–282.

Gibbs JR, van der Brug MP, Hernandez DG, Traynor BJ, Nalls MA, Lai SL, Arepalli S, Dillman A, Rafferty IP, Troncoso J et al. 2010. Abundant quantitative trait loci exist for DNA methylation and gene expression in human brain. PLoS Genet 6(5): e1000952.

Griffiths-Jones S, Grocock RJ, van Dongen S, Bateman A, Enright AJ. 2006. miRBase: microRNA sequences, targets and gene nomenclature. Nucleic acids research 34(Database issue): D140–D144.

Gu F, Doderer MS, Huang YW, Roa JC, Goodfellow PJ, Kizer EL, Huang TH, Chen Y. 2013. CMS: a web-based system for visualization and analysis of genome-wide methylation data of human cancers. PloS one 8(4): e60980.

Guo X, Gao L, Liao Q, Xiao H, Ma X, Yang X, Luo H, Zhao G, Bu D, Jiao F et al. 2013. Long non-coding RNAs function annotation: a global prediction method based on bi-colored networks. Nucleic acids research 41(2): e35.

Guzman L, Depix MS, Salinas AM, Roldan R, Aguayo F, Silva A, Vinet R. 2012. Analysis of aberrant methylation on promoter sequences of tumor suppressor genes and total DNA in sputum samples: a promising tool for early detection of COPD and lung cancer in smokers. Diagn Pathol 7: 87.

Hatada I, Fukasawa M, Kimura M, Morita S, Yamada K, Yoshikawa T, Yamanaka S, Endo C, Sakurada A, Sato M et al. 2006. Genome-wide profiling of promoter methylation in human. Oncogene 25(21): 3059–3064.

Hervouet E, Cartron PF, Jouvenot M, Delage-Mourroux R. 2013. Epigenetic regulation of estrogen signaling in breast cancer. Epigenetics : official journal of the DNA Methylation Society 8(3): 237–245.

Hill VK, Ricketts C, Bieche I, Vacher S, Gentle D, Lewis C, Maher ER, Latif F. 2011. Genome-wide DNA methylation profiling of CpG islands in breast cancer identifies novel genes associated with tumorigenicity. Cancer research 71(8): 2988–2999.

Hon GC, Hawkins RD, Caballero OL, Lo C, Lister R, Pelizzola M, Valsesia A, Ye Z, Kuan S, Edsall LE et al. 2012. Global DNA hypomethylation coupled to repressive chromatin domain formation and gene silencing in breast cancer. Genome Res 22(2): 246–258.

Hynes NE. 2000. Tyrosine kinase signalling in breast cancer. Breast Cancer Res 2(3): 154–157.

Iorio MV, Croce CM. 2012. MicroRNA dysregulation in cancer: diagnostics, monitoring and therapeutics. A comprehensive review. EMBO Mol Med 4(3): 143–159.

Irizarry RA, Ladd-Acosta C, Wen B, Wu Z, Montano C, Onyango P, Cui H, Gabo K, Rongione M, Webster M et al. 2009. The human colon cancer methylome shows similar hypo-and hypermethylation at conserved tissue-specific CpG island shores. Nat Genet 41(2): 178–186.

Jacobsen A, Silber J, Harinath G, Huse JT, Schultz N, Sander C. 2013. Analysis of microRNA-target interactions across diverse cancer types. Nat Struct Mol Biol 20(11): 1325–1332.

Jia H, Osak M, Bogu GK, Stanton LW, Johnson R, Lipovich L. 2010. Genome-wide computational identification and manual annotation of human long noncoding RNA genes. RNA 16(8): 1478–1487.

Kim JH, Dhanasekaran SM, Prensner JR, Cao X, Robinson D, Kalyana-Sundaram S, Huang C, Shankar S, Jing X, Iyer M et al. 2011. Deep sequencing reveals distinct patterns of DNA methylation in prostate cancer. Genome Res 21(7): 1028–1041.

Koga Y, Pelizzola M, Cheng E, Krauthammer M, Sznol M, Ariyan S, Narayan D, Molinaro AM, Halaban R, Weissman SM. 2009. Genome-wide screen of promoter methylation identifies novel markers in melanoma. Genome Res 19(8): 1462–1470.

Langmead B, Salzberg SL. 2012. Fast gapped-read alignment with Bowtie 2. Nature methods 9(4): 357–359.

Li D, Zhao Y, Liu C, Chen X, Qi Y, Jiang Y, Zou C, Zhang X, Liu S, Wang X et al. 2011. Analysis of MiR-195 and MiR-497 expression, regulation and role in breast cancer. Clinical cancer research : an official journal of the American Association for Cancer Research 17(7): 1722–1730.

Li X, Barkho BZ, Luo Y, Smrt RD, Santistevan NJ, Liu C, Kuwabara T, Gage FH, Zhao X. 2008. Epigenetic regulation of the stem cell mitogen Fgf-2 by Mbd1 in adult neural stem/progenitor cells. The Journal of biological chemistry 283(41): 27644–27652.

Liao Q, Liu C, Yuan X, Kang S, Miao R, Xiao H, Zhao G, Luo H, Bu D, Zhao H et al. 2011. Large-scale prediction of long non-coding RNA functions in a coding-non-coding gene co-expression network. Nucleic acids research 39(9): 3864–3878.

Lucas JJ, Domenico J, Gelfand EW. 2004. Cyclin-dependent kinase 6 inhibits proliferation of human mammary epithelial cells. Mol Cancer Res 2(2): 105–114.

Lujambio A, Calin GA, Villanueva A, Ropero S, Sanchez-Cespedes M, Blanco D, Montuenga LM, Rossi S, Nicoloso MS, Faller WJ et al. 2008. A microRNA DNA methylation signature for human cancer metastasis. Proceedings of the National Academy of Sciences of the United States of America 105(36): 13556–13561.

Ma L, Teruya-Feldstein J, Weinberg RA. 2007. Tumour invasion and metastasis initiated by microRNA-10b in breast cancer. Nature 449(7163): 682–688.

Maire V, Baldeyron C, Richardson M, Tesson B, Vincent-Salomon A, Gravier E, Marty-Prouvost B, De Koning L, Rigaill G, Dumont A et al. 2013. TTK/hMPS1 is an attractive therapeutic target for triple-negative breast cancer. PloS one 8(5): e63712.

Marsit CJ, Houseman EA, Christensen BC, Gagne L, Wrensch MR, Nelson HH, Wiemels J, Zheng S, Wiencke JK, Andrew AS, et al. 2010. Identification of methylated genes associated with aggressive bladder cancer. PloS one 5(8): e12334.

Mercer TR, Mattick JS. 2013. Structure and function of long noncoding RNAs in epigenetic regulation. Nat Struct Mol Biol 20(3): 300–307.

Nie Y, Liu X, Qu S, Song E, Zou H, Gong C. 2013. Long non-coding RNA HOTAIR is an independent prognostic marker for nasopharyngeal carcinoma progression and survival. Cancer science 104(4): 458–464.

Osman A. 2012. MicroRNAs in health and disease—basic science and clinical applications. Clin Lab 58(5-6): 393–402.

Ozsolak F, Poling LL, Wang Z, Liu H, Liu XS, Roeder RG, Zhang X, Song JS, Fisher DE. 2008. Chromatin structure analyses identify miRNA promoters. Genes & development 22(22): 3172–3183.

Pandey RR, Mondal T, Mohammad F, Enroth S, Redrup L, Komorowski J, Nagano T, Mancini-Dinardo D, Kanduri C. 2008. Kcnq1ot1 antisense noncoding RNA mediates lineage-specific transcriptional silencing through chromatin-level regulation. Mol Cell 32(2): 232–246.

Pic-Taylor A, Darieva Z, Morgan BA, Sharrocks AD. 2004. Regulation of cell cycle-specific gene expression through cyclin-dependent kinase-mediated phosphorylation of the forkhead transcription factor Fkh2p. Mol Cell Biol 24(22): 10036–10046.

Ponjavic J, Oliver PL, Lunter G, Ponting CP. 2009. Genomic and transcriptional co-localization of protein-coding and long non-coding RNA pairs in the developing brain. PLoS genetics 5(8): e1000617.

Ponting CP, Oliver PL, Reik W. 2009. Evolution and functions of long noncoding RNAs. Cell 136(4): 629–641.

Quinlan AR, Hall IM. 2010. BEDTools: a flexible suite of utilities for comparing genomic features. Bioinformatics 26(6): 841–842.

Ramsingh G, Jacoby MA, Shao J, De Jesus Pizzaro RE, Shen D, Trissal M, Getz AH, Ley TJ, Walter MJ, Link DC. 2013. Acquired copy number alterations of miRNA genes in acute myeloid leukemia are uncommon. Blood 122(15): e44–51.

Rice JC, Ozcelik H, Maxeiner P, Andrulis I, Futscher BW. 2000. Methylation of the BRCA1 promoter is associated with decreased BRCA1 mRNA levels in clinical breast cancer specimens. Carcinogenesis 21(9): 1761–1765.

Robinson MD, Stirzaker C, Statham AL, Coolen MW, Song JZ, Nair SS, Strbenac D, Speed TP, Clark SJ. 2010. Evaluation of affinity-based genome-wide DNA methylation data: effects of CpG density, amplification bias, and copy number variation. Genome Res 20(12): 1719–1729.

Rodriguez J, Frigola J, Vendrell E, Risques RA, Fraga MF, Morales C, Moreno V, Esteller M, Capella G, Ribas M et al. 2006. Chromosomal instability correlates with genome-wide DNA demethylation in human primary colorectal cancers. Cancer Res 66(17): 8462–9468.

Ruike Y, Imanaka Y, Sato F, Shimizu K, Tsujimoto G. 2010. Genome-wide analysis of aberrant methylation in human breast cancer cells using methyl-DNA immunoprecipitation combined with high-throughput sequencing. BMC genomics 11: 137.

Sanchez-Munoz A, Gallego E, de Luque V, Perez-Rivas LG, Vicioso L, Ribelles N, Lozano J, Alba E. 2010. Lack of evidence for KRAS oncogenic mutations in triple-negative breast cancer. BMC Cancer 10: 136.

Shen J, Wang S, Zhang YJ, Kappil MA, Chen Wu H, Kibriya MG, Wang Q, Jasmine F, Ahsan H, Lee PH et al. 2012. Genome-wide aberrant DNA methylation of microRNA host genes in hepatocellular carcinoma. Epigenetics : official journal of the DNA Methylation Society 7(11): 1230–1237.

Suzuki H, Maruyama R, Yamamoto E, Kai M. 2012. DNA methylation and microRNA dysregulation in cancer. Mol Oncol 6(6): 567–578.

Trapnell C, Williams BA, Pertea G, Mortazavi A, Kwan G, van Baren MJ, Salzberg SL, Wold BJ, Pachter L. 2010. Transcript assembly and quantification by RNA-Seq reveals unannotated transcripts and isoform switching during cell differentiation. Nat Biotechnol 28(5): 511–515.

Vrba L, Munoz-Rodriguez JL, Stampfer MR, Futscher BW. 2013. miRNA gene promoters are frequent targets of aberrant DNA methylation in human breast cancer. PloS one 8(1): e54398.

Wamstad JA, Alexander JM, Truty RM, Shrikumar A, Li F, Eilertson KE, Ding H, Wylie JN, Pico AR, Capra JA et al. 2012. Dynamic and coordinated epigenetic regulation of developmental transitions in the cardiac lineage. Cell 151(1): 206–220.

Wang C, Bian Z, Wei D, Zhang JG. 2011. miR-29b regulates migration of human breast cancer cells. Molecular and cellular biochemistry 352(1-2): 197–207.

Weber B, Stresemann C, Brueckner B, Lyko F. 2007. Methylation of human microRNA genes in normal and neoplastic cells. Cell Cycle 6(9): 1001–1005.

Wu WK, Lee CW, Cho CH, Fan D, Wu K, Yu J, Sung JJ. 2010. MicroRNA dysregulation in gastric cancer: a new player enters the game. Oncogene 29(43): 5761–5771.

Xu H, Wei CL, Lin F, Sung WK. 2008. An HMM approach to genome-wide identification of differential histone modification sites from ChIP-seq data. Bioinformatics 24(20): 2344–2349.

Yan LX, Huang XF, Shao Q, Huang MY, Deng L, Wu QL, Zeng YX, Shao JY. 2008. MicroRNA miR-21 overexpression in human breast cancer is associated with advanced clinical stage, lymph node metastasis and patient poor prognosis. RNA 14(11): 2348–2360.

Zhang B, Kirov S, Snoddy J. 2005. WebGestalt: an integrated system for exploring gene sets in various biological contexts. Nucleic acids research 33(Web Server issue): W741–748.

Zhang L, Sullivan PS, Goodman JC, Gunaratne PH, Marchetti D. 2011a. MicroRNA-1258 suppresses breast cancer brain metastasis by targeting heparanase. Cancer research 71(3): 645–654.

Zhang Z, Zhang B, Li W, Fu L, Zhu Z, Dong JT. 2011b. Epigenetic Silencing of miR-203 Upregulates SNAI2 and Contributes to the Invasiveness of Malignant Breast Cancer Cells. Genes & cancer 2(8): 782–791.

